# Loss of an uncharacterized mitochondrial methionine tRNA-synthetase induces mitochondrial unfolded protein response in *Caenorhabditis elegans*

**DOI:** 10.1101/2025.02.03.636310

**Authors:** Bharat Vivan Thapa, Mohit Das, James P Held, Maulik R Patel

## Abstract

Aminoacyl-tRNA synthetases (aaRSs) are essential for translation, as they charge tRNA molecules with their corresponding amino acids. Alterations in aaRSs can significantly disrupt both cytosolic and mitochondrial translation. Through a forward genetic screen for mitochondrial unfolded protein response (UPR^mt^) activators in *C. elegans*, we identified a missense mutation (P447V) in the previously uncharacterized gene Y105E8A.20, which encodes a dually localized methionine tRNA synthetase (MetRS). Here, we characterize the UPR^mt^ induction by Y105E8A.20, which we call *mars-2*, and demonstrate that the P447V allele is a loss-of-function mutation. Furthermore, we show impaired *mars-2* activity in the mitochondria triggers UPR^mt^. This strain provides a valuable tool for studying mitochondrial translation and understanding how aaRSs are involved in mitochondrial homeostasis.

## Introduction

Mitochondria are multifaceted organelles involved in a wide range of essential processes, including energy production, intracellular signaling, and programmed cell death^1,2^. This semi-autonomous organelle has its own genome. In *Caenorhabditis elegans*, the mitochondrial genome encodes 36 genes, including essential protein subunits of the mitochondrial respiratory complexes, mitochondrial rRNAs, and mitochondrial tRNAs^3^. The mitochondrial genome is transcribed as a single polycistronic RNA, which is subsequently processed and then mRNAs are translated within mitochondria by specialized mito-ribosomes^4^. While mitochondrial tRNAs are encoded within the mitochondria, they must be charged with their corresponding amino acids to locally translate mitochondrial DNA encoded proteins. This charging process is carried out by aminoacyl tRNA synthetases (aaRSs), which are encoded by the nuclear genome and imported into the mitochondria^5^. aaRSs are highly specialized enzymes involved in covalently attaching amino acids to their cognate tRNAs. Given their house-keeping function across eukaryotes, bacteria, and archaea^6,7^, mutations in aaRS genes are linked to Charcot-Marie-Tooth (CMT) disease^8–12^, tumor formation^13–15^, and death^16^. Recent studies show that mutations in aaRS can alter tRNA levels, which in turn regulate the production of tRNA-derived fragments and contribute to diverse disease phenotypes^17^.

Although mitochondrial aaRS mutations are associated with several diseases^18–20^, the mechanisms by which these mutations affect mitochondrial function and contribute to pathological phenotypes remains largely unknown. Dysfunctional aaRS activity can directly affect mitochondrial protein synthesis causing mitochondrial stress^21,22^. Mitochondrial quality control pathways, including the mitochondrial unfolded protein response (UPR^mt^) are activated in response to mitochondrial stress^23^. UPR^mt^ upregulates nuclear-encoded chaperones and proteases to preserve mitochondrial functionality^24,25^. In *C. elegans*, this response is mediated by the transcription factor ATFS-1, which in healthy cells is imported to the mitochondria^26^. However when mitochondria are stressed, a decrease in mitochondrial protein import disrupts ATFS-1 mitochondrial import^27^, resulting in ATFS-1 nuclear accumulation and subsequent transcription of UPR^mt^-associated genes. Transcriptional reporters, such as the *hsp-6p::GFP* strain (one of the UPR^mt^-associated genes), have been developed to monitor UPR^mt^ activation^28^.

This study reveals a compelling link between mitochondrial protein translation fidelity and the activation of the mitochondrial unfolded protein response. We demonstrate a missense mutation in a previously uncharacterized mitochondrial methionine tRNA synthetase gene in *C. elegans* that activates UPR^mt^. Using CRISPR-Cas9, we confirmed the causality of this point mutation and demonstrated that it represents a loss-of-function mutation. Furthermore, we mutated the mitochondrial targeting sequence (MTS) and found that the absence of MARS-2 in the mitochondria is sufficient to induce UPR^mt^. These findings not only enhance our understanding of mitochondrial quality control but also identify MARS-2 as a potential target for developing therapeutic strategies to modulate UPR^mt^ in conditions involving mitochondrial dysfunction.

## Results

### P447 is a highly conserved amino acid residue in MARS-2

In a previous study^29^, we used *hsp-6p::GFP* strain (Figure 1A) to perform a forward genetic screen aimed at identifying inducers of UPR^mt^ (Figure 1B). This screen uncovered a missense mutation (P447V) in the gene *Y105E8A.20*, which encodes a methionine tRNA synthetase in *C. elegans* (mpt139 strain shown in Figure 1C and 1D) along with other non-causal mutations introduced through EMS mutagenesis. Phylogenetic analysis of protein sequences from cytoplasmic and mitochondrial methionine tRNA synthetases (MARS1 and MARS2, respectively) across various species revealed that Y105E8A.20 protein in *C. elegans* – hereafter referred as MARS-2 – shares greater similarity with MARS-2 proteins compared to MARS-1 proteins in other species (Figure 1E). Furthermore, protein sequence alignment using CLUSTAL tool established that the 447^th^ amino acid residue is highly conserved across species (Figure 1F).

**Figure 1:**
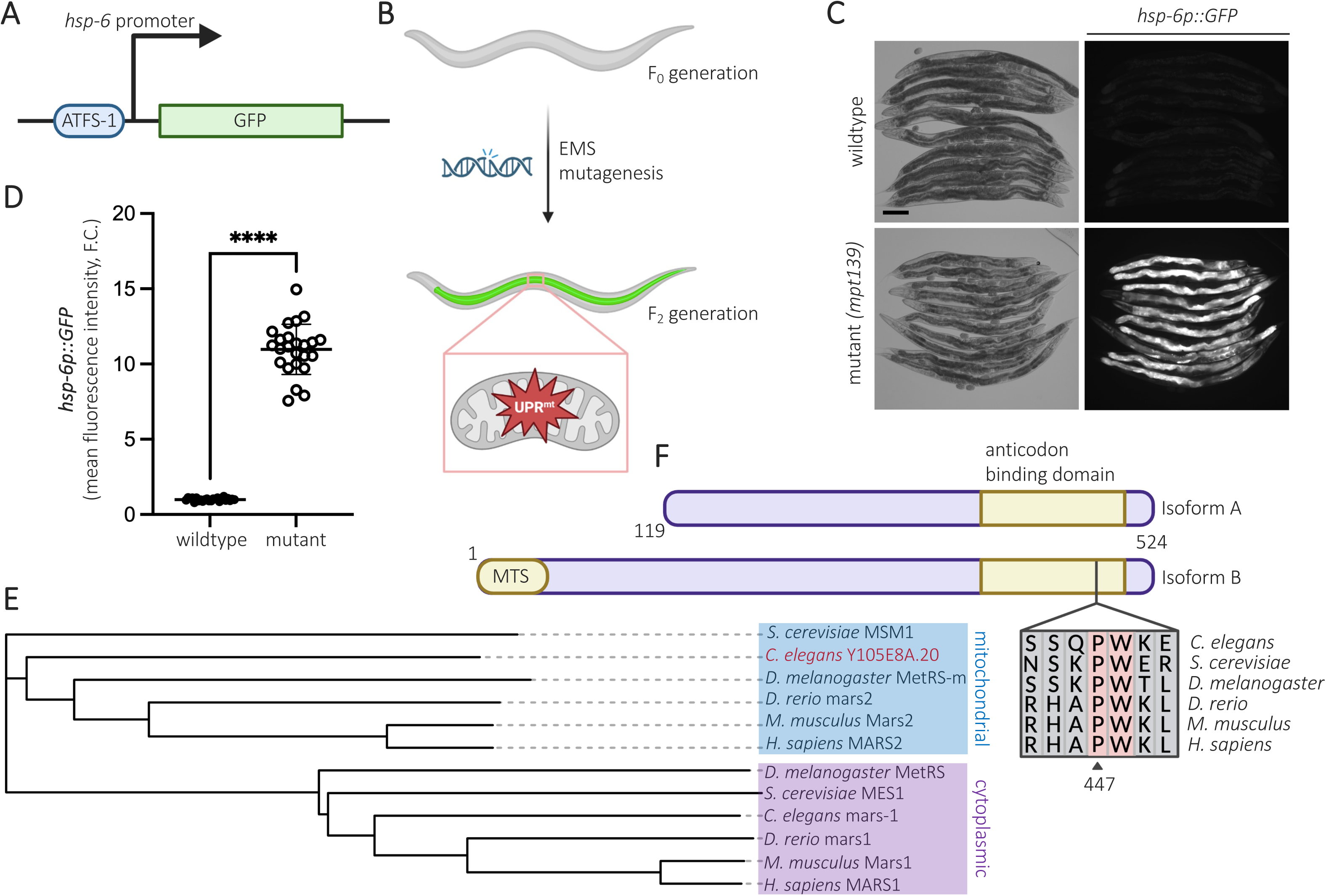
Phylogenetic analysis show that P447 is a highly conserved amino acid residue across species. (A) Gene transcription diagram depicting the expression of *hsp-6p::GFP* transcriptional reporter. (B) Schematic for the forward genetic screen to identify activators of mitochondrial unfolded protein response. (C) Representative brightfield and corresponding fluorescent micrographs showing UPR^mt^ reporter (*hsp-6p::GFP*) activation in day 2 (D2) adult wildtype and mutant (*mpt139)* animals. (D) Scatter plot showing fluorescence intensity quantification of the UPR^mt^ reporter in wildtype and mutant animals normalized to the reporter intensity in wildtype animals (n = 24, mean and SD shown, unpaired t-test with Welch’s correction). (E) Phylogenetic tree illustrating evolutionary relationships between species based on their mitochondrial and cytoplasmic methionine tRNA synthetase protein sequences (Y105E8A.20 highlighted in red). (F) Pictorial depiction of the two isoforms of MARS-2 protein, along with the conserved proline residue (P447) in the anticodon binding domain of the protein in *C. elegans*. Scale bar – 200 µm, GFP – green fluorescent protein, MTS – mitochondrial targeting sequence, F.C. – Fold change, ns – non-significant, * – P < 0.05, ** – P < 0.01, *** – P < 0.001, **** – P < 0.0001

### P447V mutation in MARS-2 induces mitochondrial unfolded protein response

To confirm the role of the P447V mutation in activating UPR^mt^, we introduced this missense mutation into the *zcIs13 V* background (*hsp-6p::GFP* strain) using CRISPR-Cas9 gene editing. Two independent clones carrying the desired mutation were recovered and validated through Sanger sequencing. Fluorescent imaging of whole animal revealed that the P447V mutant exhibited a 7-fold increase in UPR^mt^ reporter activation compared to wildtype animals (Figure 2A and 2B), confirming the mutation’s causative role in inducing UPR^mt^.

**Figure 2:**
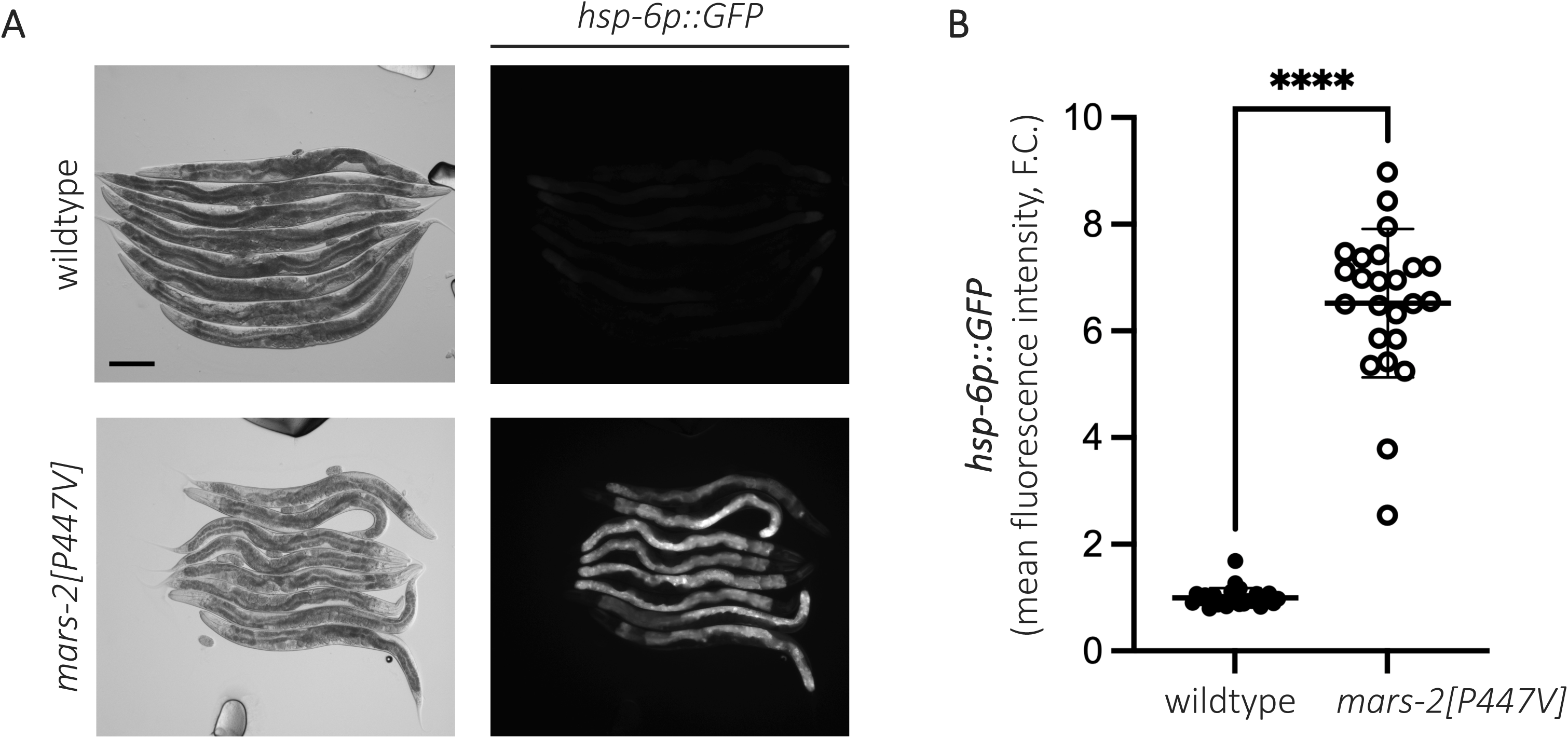
P447V is the causal mutation in MARS-2 that induces mitochondrial unfolded protein response. (A) Representative brightfield and corresponding fluorescent micrographs showing UPR^mt^ reporter (*hsp-6p::GFP*) activation in day 2 (D2) adult wildtype and *mars-2[P447V]* animals. (C) Scatter plot showing fluorescence intensity quantification of the UPR^mt^ reporter in wildtype and *mars-2[P447V]* animals normalized to the reporter intensity in wildtype animals (n = 24, mean and SD shown, unpaired t-test with Welch’s correction). Scale bar – 200 µm.

### P447V mutation in MARS-2 induces mitochondrial unfolded protein response via ATFS-1 signaling

Previous studies have shown that ATFS-1 plays a central role in inducing UPR^mt^ in *C. elegans*^26,27^. To determine whether the P447V mutation in MARS-2 activates UPR^mt^ through this pathway, we performed RNAi-mediated knockdown of *atfs-1* in the mutant strains. This completely suppressed reporter activation (Figure 3A and 3B), confirming that UPR^mt^ activation in the mutant strains occurs via the canonical ATFS-1 pathway. Interestingly, animals harboring the P447V mutation arrested upon *atfs-1* knockdown (Figure 3A). This is likely due to their inability to mount a sufficient stress response in the absence of ATFS-1, consistent with other reports that ATFS-1 is protective against multiple stressors^30^. Collectively, these findings establish the P447V mutation in *mars-2* gene as a driver of UPR^mt^ activation via ATFS-1.

**Figure 3:**
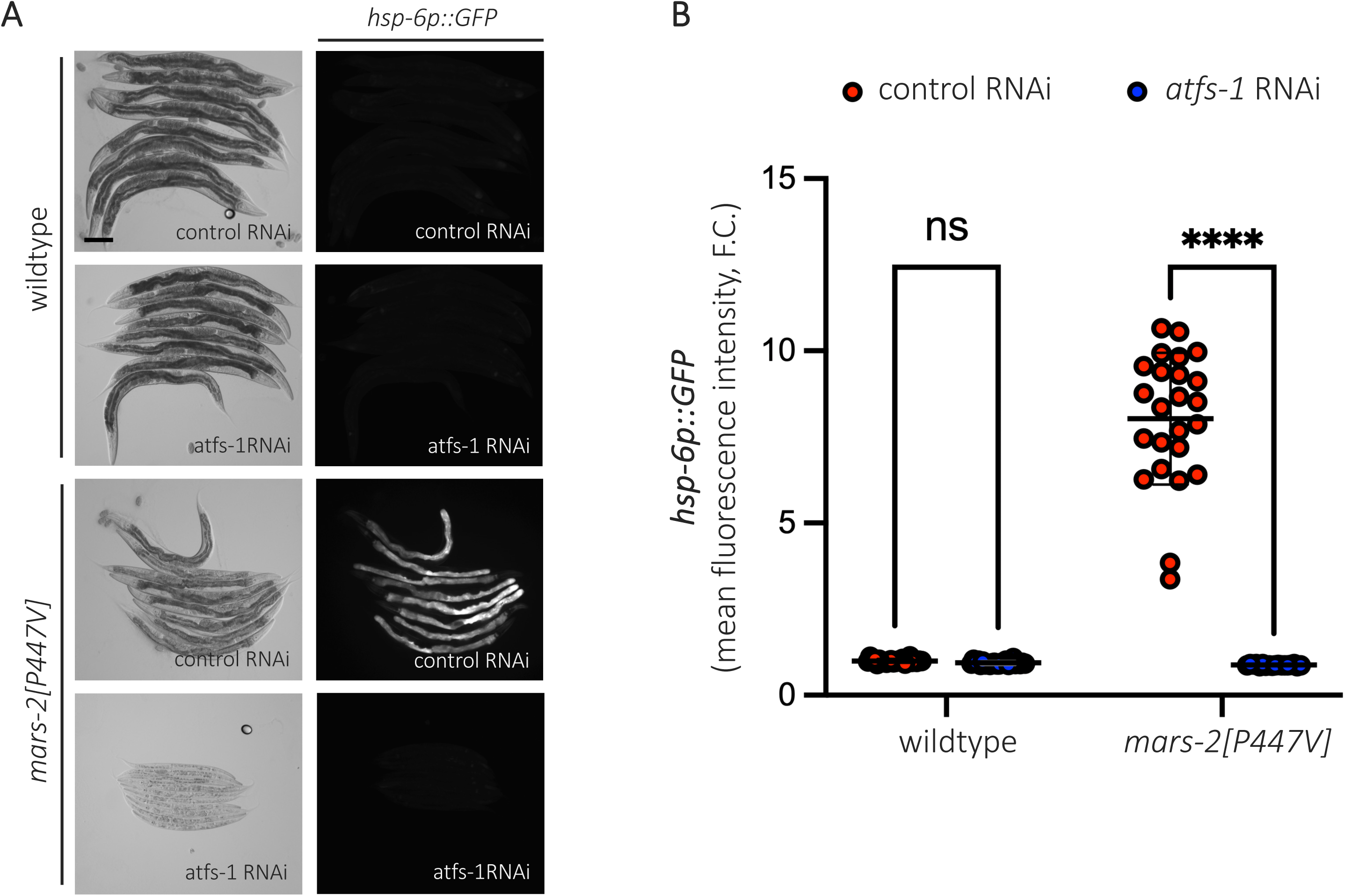
MARS-2(P447V)-mediated mitochondrial unfolded protein response is ATFS-1 dependent. (A) Representative brightfield and corresponding fluorescent micrographs showing UPR^mt^ reporter activation in D2 adult wildtype and *mars-2[P447V]* animals upon control and *atfs-1* RNAi. (B) Scatter plot showing fluorescence intensity quantification of the UPR^mt^ reporter in wildtype and *mars-2[P447V]* animals upon control and *atfs-1* RNAi normalized to the reporter intensity in wildtype animals on control RNAi (n = 18 in *mars-2[P447V]* on *atfs-1* RNAi, n=24 others, mean and SD shown, two-way ANOVA with Sidak’s multiple comparison test). Scale bar – 200 µm.

### P447V mutation in MARS-2 is a loss-of-function mutation

Next, we asked what effect this mutation has on the protein. InterPro domain analysis identified the P447V mutation within the anticodon-binding domain of the MARS-2 protein, a region critical for recognizing the anticodon loop of methionine tRNA. Given that mutations in the anticodon-binding domain of other aaRSs are often associated with reduced enzymatic activity^19,31^, we hypothesized that P447V mutation in *mars-2* represents a loss-of-function mutation. To test this, we performed RNAi targeting Y105E8A.20 transcripts in wildtype and mutant strains. RNAi in wildtype animals induced UPR^mt^ (Figure 4A and 4B), mimicking the effect of the P447V mutation. Furthermore, RNAi in P447V mutant animals resulted in an even greater activation of UPR^mt^ and exacerbated growth defects (Figure 4A and 4B), consistent with P447V being a loss-of-function mutation.

**Figure 4:**
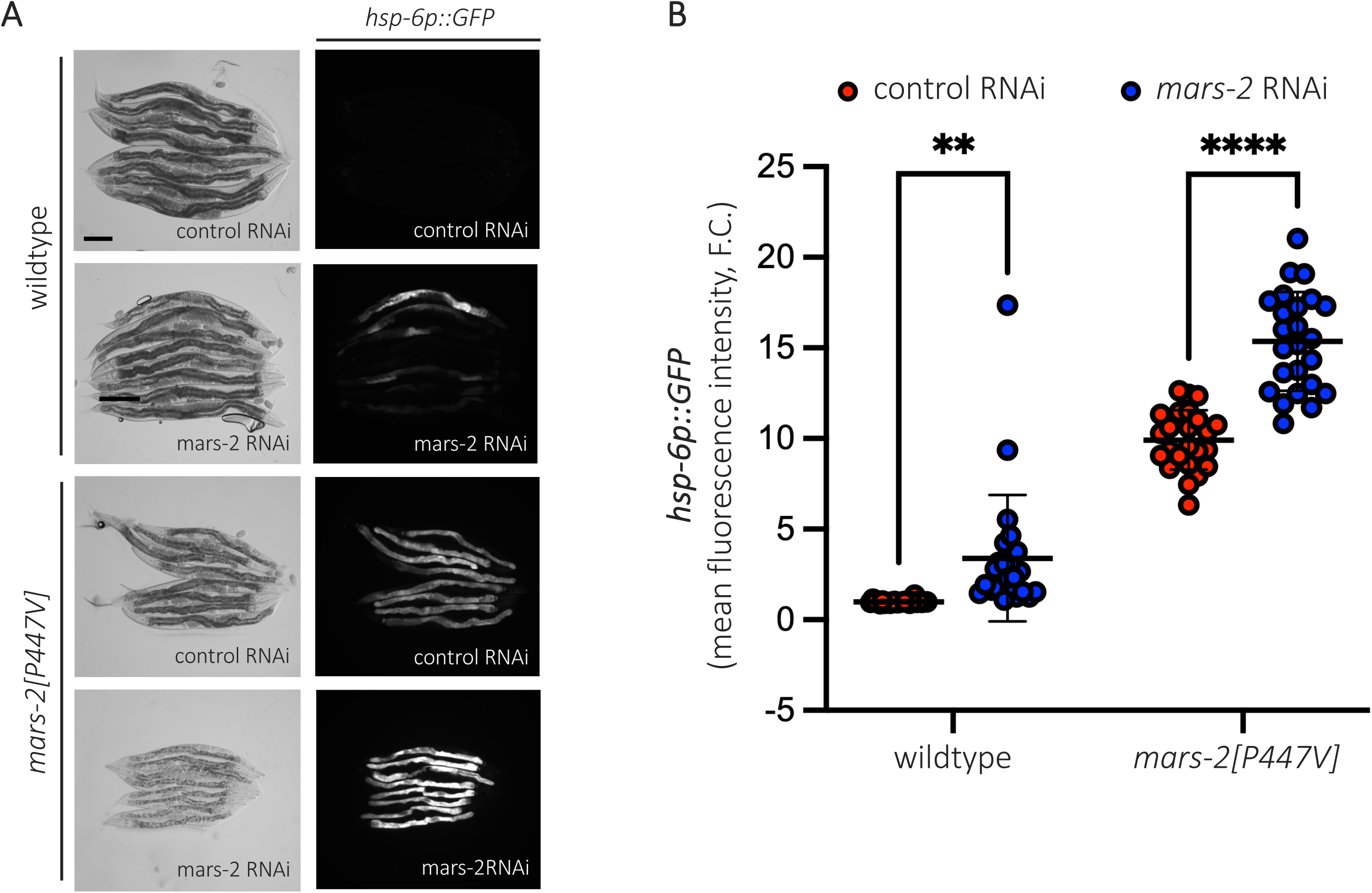
P447V mutation in MARS-2 is a loss-of-function mutation. (A) Representative brightfield and corresponding fluorescent micrographs showing UPR^mt^ reporter activation in D2 adult wildtype and *mars-2[P447V]* animals upon control and *mars-2* RNAi. (G) Scatter plot showing fluorescence intensity quantification of the UPR^mt^ reporter in wildtype and *mars-2[P447V]* animals upon control and *mars-2* RNAi normalized to the reporter intensity in wildtype animals on control RNAi (n = 24, mean and SD shown, two-way ANOVA with Sidak’s multiple comparison test). Scale bar – 200 µm.

### Mitochondrial loss of MARS-2 activates robust mitochondrial unfolded protein response

The *mars-*2 gene encodes for two isoforms: isoform A (406 aa) and isoform B (524 aa). Analysis using MitoFates software predicts a mitochondrial targeting sequence (MTS) within the N-terminal region of isoform B, absent in isoform A. To determine whether loss of *mars-2* mitochondrial function specifically underpins UPR^mt^ activation, we generated a mitochondrial-specific loss-of-function mutant, *mpt227* by replacing the first methionine with alanine using CRISPR-Cas9 gene editing. This results in mutants that lack the mitochondrial isoform but retained the shorter, likely cytoplasmic isoform. Consistent with a previous study where RNAi knockdown of several mitochondrial ARS genes induced UPR^mt^ ^32^, we found that the mitochondrial-specific loss-of-function mutants (labelled as *mars-2[*Δ*MTS]*) exhibited robust UPR^mt^ (Figure 5A and 5B). Moreover, we concluded that P447V is likely a hypomorphic mutation in *mars-*2, which compromises its mitochondrial function but remains a viable strain, in contrast to the severe growth defects and developmental arrest observed in the mitochondrial-specific loss-of-function mutants (Figure 5A and 5C). These findings suggest that mitochondrial isoform of *mars-2* is essential in maintaining mitochondrial homeostasis.

**Figure 5:**
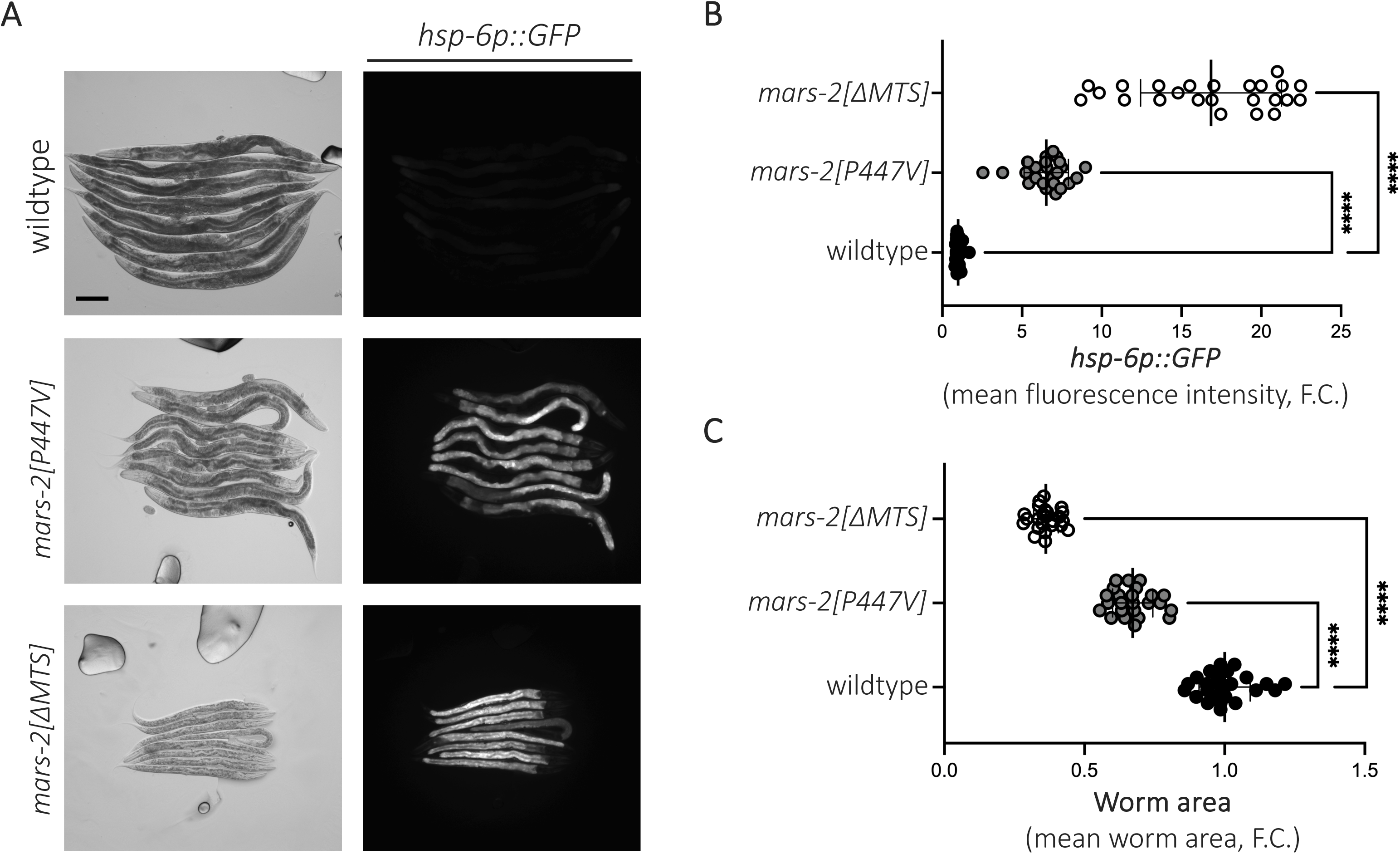
Loss of mitochondrial MARS-2 isoform by disrupting MTS activates mitochondrial unfolded protein response. (A) Representative brightfield and corresponding fluorescent micrographs showing UPR^mt^ reporter activation in D2 adult wildtype, *mars-2[P447V],* and *mars-2[*Δ*MTS]* animals. (B) Scatter plot showing fluorescence intensity quantification of the UPR^mt^ reporter in wildtype, *mars-2[*P447V], and *mars-2[*Δ*MTS]* animals normalized to the reporter intensity in wildtype animals (n = 24, mean and SD shown, one-way ANOVA with Dunnett’s multiple comparison test) (C) Scatter plot depicting outline area quantification of wildtype, *mars-2[P447V]*, and *mars-2[*Δ*MTS]* animals (n = 24, mean and SD shown, one-way ANOVA with Dunnett’s multiple comparison test). Scale bar – 200 µm.

## Discussion

Our study demonstrates that the P447V mutation in the *mars-2* gene is a loss-of-function mutation that induces UPR^mt^. One reasonable possibility is that the mutation disrupts the function of mitochondrial methionine-tRNA synthetase. Supporting this, we found that preventing MARS-2 from entering mitochondria also strongly activates UPR^mt^ (Figure 5A and 5B). As the P447V mutation occurs in a highly conserved anticodon binding domain of the enzyme (Figure 1F), we hypothesize it reduces methionine tRNA charging, thereby affecting mitochondrial protein production. Given that defects in mitochondrial translation are known to trigger UPR^mt^ ^33,34^, P447V mutation in *mars-2* may induce UPR^mt^ by disrupting mitochondrial protein production. In the future, this can be further tested by measuring charged methionine levels and assessing mitochondrial protein production in P447V mutants. Additionally, our phylogenetic analysis of MARS-2 in *C. elegans* reveals functional similarity with mitochondrial methionine tRNA-synthetases from other species (Figure 1E).

In this study, we did not investigate the effects of P447V mutation on the cytoplasmic isoform of *mars-2* in *C. elegans*. Considering the functional role of *let-65* (*mars-1*) as the cytoplasmic methionyl-tRNA synthetase^35^, it is plausible that any deleterious effects of P447V mutation in the cytoplasmic isoform of *mars-2* are mitigated due to functional redundancy. Interestingly, UPR^mt^ activation in P447V mutants was restricted to the intestine (Figure 1C, 2A, 3A, 4A, and 5A). This tissue-specific response parallels findings in humans, where mitochondrial aaRS mutations predominantly affect high-energy-demand tissues such as muscles and the nervous system^36^. The intestinal specificity observed in these mutants may reflect intrinsic differences in mitochondrial dynamics or stress sensitivity between tissues.

While this study establishes a strong link between the P447V mutation in *mars-2* and UPR^mt^ activation, the precise mechanism by which mitochondrial MetRS dysfunction triggers UPR^mt^ remain to be explored. The mutant strains developed in this study provide a valuable tool for mitochondrial biology and the pathological consequences of aaRS mutations.

## Materials and methods

### Worm strains

All the strains used in this study are listed in the table below:

| Genotype | Source | Strain name |
| --- | --- | --- |
| *C. elegans* wildtype | CGC | N2 |
| *zcls13[hsp-6p::GFP] V* | CGC | GL347 |
| *mptsi1, zcls13 V; mpt139* | Held et al<br>(2024)<sup>29</sup> | MRP649 |
| *mars-2(mpt213[P447V]) I* | This study | MRP1019 |
| *mars-2(mpt215[P447V]) I; zcls13 V* | This study | MRP1021 |
| *mars-2(mpt216[P447V]) I; zcls13 V* | This study | MRP1022 |
| *mars-2(mpt227[ΔMTS]) I +/-; zcls13 V* | This study | MRP1042 |
| *mars-2(mpt228[ΔMTS]) I +/-; zcls13 V* | This study | MRP1043 |

### Worm maintenance

All worms were raised at 20 °C on nematode growth media (NGM) seeded with OP50 *Escherichia coli* bacteria.

### CRISPR/Cas9 gene editing

CRISPR was performed as previously described^37^ using Alt-R S.p. Cas9 Nuclease V3 (IDT #1081058), tracrRNA (IDT #1072532) and using *dpy-10* as a co-injection marker^38^. The crRNA sequence used for introducing P447V mutation in *mars-2* was TCAATCAAGTCAACCATGGA and ssODN sequence: TGAAAGAAGCGAATCGATTATTTCAATCAAGTCAATCATGGAAAGAAATTGATGAAAAACGGCTGAAAAGTCTACTTTT. The crRNA sequence used for introducing ΔMTS mutation in *mars-2* was 5’ – CGCTTTTCATGAATCCTTGG – 3’ and ssODN sequence: 5’ – cgattgattaattcacttttttttttgcgcttttcGCTAATCCATGGAGATTTTTCGTGAGAAAATCGAGTACATTTGTCACTTC– 3’. The Cas9::tracrRNA::crRNA complexes and repair templates were microinjected into the zcIs13 strain to obtain *mpt215* and *mpt227*, respectively. The *dpy-10* co-injection marker was then outcrossed using a wildtype (N2, RRID: WBStrain00000001) strain.

### Genetic crosses

The genetic crosses were performed by crossing 5 larval stage 4 (L4) hermaphrodites of a strain to 15 heterozygous males of another strain (these were first obtained by crossing L4 hermaphrodites of the strain with wildtype males). An appropriate number of F1 generation L4 hermaphrodites were cloned out of the cross plate and allowed to have self-progeny. An appropriate number of F2 progeny were cloned and genotyped for allele of interest once they had progeny. The genotyping was performed using IDT primers and NEB enzymes below:

For genotyping P447V mutation in *mars-2,* Forward – GGAAATGGTCGAGGAGAGCAGAG Reverse –CCTCCAGAATAGCTTCCAAATCGGG and NcoI cuts wildtype amplicon only. For genotyping ΔMTS mutation in *mars-2,* Forward – CAGGAATTGAAGGAGCTAAATTCTGCAAAG Reverse – CGAAATTAGGGATTATCAGACGCAAGTTCC and NcoI-HF cuts mutant amplicon only.

### Fluorescence microscopy

Zeiss Axio Zoom V16 stereo zoom microscope was used for imaging fluorescent UPR^mt^ reporter in whole animals. The worms were mounted on 2% agarose pads on microscopic slides and immobilized using 1 µL of 100 mM levamisole (ThermoFisher #AC187870100). A coverslip was used to avoid the worms from drying.

### Image analysis

The brightfield and fluorescent animal images were analyzed using the Fiji application. The outline of each worm was traced as the region of interest on the brightfield snapshot. The ‘Measure’ feature under ‘Analyze’ tab was used for measuring the area of the animal and its fluorescence intensity (calculated by summing all the pixel values within the region of interest and dividing by the total number of pixels in that area) in the fluorescence snapshot.

### RNA interference

RNAi was performed by shifting embryos to NGM plates seeded with RNAi bacteria, as described previously^39^. *E. coli* HT115 carrying specific RNAi clones were grown overnight from a single colony in 2 mL of LB supplemented with 50 μg/mL ampicillin. To make 16 RNAi plates, 50 ml of LB supplemented with 50 μg/ml ampicillin and inoculated with 500 μL of the overnight culture, then incubated while shaking at 37 °C for 4–5 hours (to an OD_550-600_ of ∼0.8). Next, to induce expression of the double stranded RNA, cultures were supplemented with an additional 50 ml of LB supplemented with 50 μg/ml ampicillin and 4 mM IPTG and then continued to incubate while shaking at 37 °C for 4 hours. Following incubation, bacteria were pelleted by centrifugation at 3900 rpm for 6 min. Supernatant was decanted and pellets were gently resuspended in 4 ml of LB supplemented with 8 mM IPTG. 250 μl of resuspension was seeded onto standard NGM plates containing 1 mM IPTG. Plates were left to dry overnight and then used within 2 weeks. Bacterial RNAi strains were from Ahringer RNAi Feeding Library, grown from single colony and identity confirmed by Sanger sequencing. *Y105E8A.20* (Y105E8A.20), *atfs-1* (ZC376.7).

### Statistical analysis

For whole animal fluorescence analysis, a sample size of 24 was used in the study and each animal was considered a biological replicate. The p-values under 0.05 were considered significant while p-values above 0.05 were considered non-significant (ns). All statistical analysis was performed using Prism 10 and the information on specific statistical test can be found in the figure legend.

## Data availability

Strains are available upon request. The authors affirm that all data necessary for confirming the conclusions of the article are present within the article and figures.

## Acknowledgements

We are grateful to WormBase for providing valuable information that supported our research on *mars-2* in *C. elegans* (Okimoto et al., 1992). Protein alignment for various homologs of the MARS2 protein was conducted using CLUSTAL Omega, a multiple sequence alignment tool developed by EMBL and phylogenetic tree was constructed using iTOL (Interactive Tree Of Life). The visual representation of *mars-2* was created using BioRender. Image analysis was performed using Fiji, and graphs were generated with GraphPad Prism.

## Author contributions

Bharat Vivan Thapa – Writing - original draft, Investigation, Methodology, Validation, Formal analysis, Visualization, Conceptualization

Mohit Das – Investigation, Methodology, Validation, Formal analysis, Visualization, Conceptualization

James P. Held – Conceptualization, Investigation, Methodology, Writing - review & editing Maulik R. Patel – Supervision, Writing - review & editing, Funding acquisition, Project administration, Conceptualization

